# Axonopathy in Duchenne Muscular Dystrophy limits microdystrophin gene therapy efficacy

**DOI:** 10.64898/2026.08.17.744857

**Authors:** Aly Bourguiba-Villeneuve, Maxime Gelin, Aurélie Fail, Lucile Saillard, Stéphanie Bauché, Cécile Peccate, Pierre Meunier, Zoheir Guesmia, Lou Ann Mirabile, Julianne Perronnet, Mégane Lemaitre, Lorenzo Giordani, Sestina Falcone, Christel Gentil, France Pietri-Rouxel

## Abstract

Duchenne muscular dystrophy (DMD) is classically defined as a primary myopathy, and current AAV-mediated microdystrophin gene therapies are shown to successfully preserve muscle integrity. However, their efficacy in recovering functional outcomes remains to improve. We hypothesized that this limitation stems from an unaccounted vulnerability within the peripheral nerve. Here, we demonstrate that the mdx mouse model exhibits a peripheral axonopathy independently of muscle necrosis. Using single-nucleus RNA sequencing and structural analyses, we have identified an active denervation program and a profound failure of neural repair pathways. Importantly, we revealed that the full-length dystrophin isoform Dp427c is expressed in the healthy peripheral nerve, intimately following the cytoskeletal organization and accumulating at regions of high biomechanical stress, including Schmidt-Lanterman incisures and Nodes of Ranvier. In its absence, nerves of mdx mice loss an essential scaffolding support, leading to localized structural collapse. Furthermore, we showed that muscle-restricted microdystrophin gene therapy rescues sarcolemmal integrity but failed to restore nerve-muscle connectivity or resolved neurotransmission defects. These findings fundamentally redefine DMD as an integrated motor unit pathology, thereby underscoring the absolute necessity of implementing combined therapeutic strategies that target both the muscle and the peripheral nervous system.

## INTRODUCTION

Duchenne muscular dystrophy (DMD) is a devastating neuromuscular disorder caused by mutation in the DMD gene, resulting in the absence of the dystrophin protein(1). The loss of dystrophin leads to the destabilization of the dystrophin-glycoprotein complex (DGC), which anchors the actin cytoskeleton to the plasma membrane(1). This instability causes progressive muscle fiber degeneration, respiratory failure and cardiomyopathy(2).

Among the therapeutic approaches currently in clinical development, adeno-associated virus (AAV)- mediated gene therapy is the most promising(3–7). In particular, AAV serotypes -8 or 9 are used for their efficacy to transduce muscle fibers, including diaphragm and heart(8). Due to the packaging limitation of these viral vectors, a truncated dystrophin, microdystrophin, containing the essential functional domains of the protein under the control of a muscle-specific promoter, has been developed(5,9). However, despite successful AAV delivery and the consequent improvement of muscle damage, measured by decrease of Creatine Kinase (CK) level in blood serum(10), the efficacy of these therapy on functional parameters are still debated or even unconvincing despite of administration of high-dose of AAV, reported to induce significant toxicity and serious side effects(7,11).

Several studies have established that the neuromuscular junction (NMJ) exhibits severe morphological and functional defects in DMD(12,13). The DGC is not only essential for the clustering of acetylcholine receptors (AChRs) but has also been described as implicated in axonal guidance. Consequently, its absence severely disrupts both the structural organization and signal transmission at the motor endplate(12). Since DMD disease is canonically defined as a myopathy, these neural defects have been regarded as mere secondary consequences of myofiber necrosis, in accordance with a passive “dying-back” hypothesis. Crucially, the existing literature only observes this structural damage without elucidating its cause(14,15). Indeed, descriptive investigations in the mdx mouse model have reported peripheral nerve alterations, including demyelination and a reduction in large-diameter axons without deciphering the molecular origin(14). Up to date, it remains unknown whether the observed axonopathy is truly a secondary collateral effect, or if the peripheral nerve possesses an intrinsic requirement for dystrophin, making its degeneration an autonomous event.

With the ambition of further optimizing current AAV-microdystrophin approaches, which have already demonstrated remarkable success in preserving muscle integrity(7,10,11,16), this study investigated the overlooked neural compartment of the motor unit. We hypothesized that the limited functional outcomes observed with AAV-microdystrophin treatment might stem from an unaccounted-for vulnerability in the peripheral nerve itself. To test this, we first evidenced the presence of full-length dystrophin within the healthy sciatic nerve and characterized the autonomous axonopathy resulting from its absence in the mdx model. By subsequently evaluating a muscle-restricted microdystrophin therapy, we confirmed its efficacy in fully rescuing muscle contractile capacity. However, we demonstrated that repairing muscle sarcolemmal integrity alone is not sufficient to fully re-establish nerve-muscle connectivity or resolve neurotransmission defects. Ultimately, this work sheds light on the mechanisms at play in order to further improve current clinical strategies, emphasizing that supporting the motor unit as a whole is the next crucial step in maximizing the functional outcomes of gene therapies for DMD.

## RESULTS

### Pathological muscle hypertrophy and neurogenic atrophy follow a biphasic longitudinal course

To decode the complex temporal dynamics governing dystrophic muscle degeneration, we evaluated biometric, histological and molecular courses in mdx and wild-type (WT) mice across a chronological continuum from 9 to 62 weeks of age (Fig 1A). Systemic biometric monitoring revealed that the total body weight (BW) remained comparable between WT and mdx up to 62 weeks of age, after which mdx exhibited a significant increase in overall BW compared to age-matched WT mice (Fig 1B). Concurrently, morphometric evaluation of the Tibialis anterior (TA) muscle mass (Fig S1A) normalized to BW unveiled a distinct biphasic kinetic profile. Indeed, in early adulthood (9 to 14W), mdx mice displayed a marked pathological hypertrophy of the TA, with the TA/BW ratio peaking at 14W and beyond this 14W threshold, the TA mass decline (Fig.1C). At the cellular level, analysis of myofiber cross-sectional area (CSA) confirmed this decrease in muscle mass. In contrast to the stable morphological profile of TA in WT, the mdx muscle exhibited a progressive shift toward an atrophic state, characterized by an accumulation of small-caliber fibers (250-1000µm^2^) beginning at 14W (Fig S1B). Given that chronic denervation is a well-established driver of neurogenic muscle atrophy, we hypothesized that a loss of synaptic connectivity might induce this muscle mass decline. To elucidate theses underlying mechanisms, we systematically evaluated neuromuscular junction (NMJ) remodeling and synaptic stability markers across the time course. At each time point analyzed, quantitative transcriptional profiling revealed a significant and sustained upregulation of the acetylcholine receptor (AChR) alpha-1 (*Chrna1*) and the embryonic gamma (*Chrng*) subunits in mdx muscles (Fig 1D-E). This chronic induction of AChR subunit expression pointed out ongoing synaptic instability and compromised neuromuscular transmission. In parallel, a striking downregulation of the transcript of the Schwann cells (SCs)-specific marker *S100b*, occurred across all time points, demonstrating a continuous and critical alteration of glial support at the NMJ (Fig 1F). These molecular signatures of chronic synaptic distress were substantiated by structural and spatial alterations at the tissue level. High-resolution topological mapping of AChR distribution on muscle cross-sections revealed that while WT mice maintained strictly localized and compact AChR clusters, mdx muscle exhibited a widespread extra-synaptic dispersion of AChR clusters across the entire muscle section at all evaluated timepoints (Fig 1G). In addition, we analyzed the spatial rearrangement of individual myofibers to assess the ongoing remodeling of motor units. Furthermore, unlike WT muscles, which maintained a classically described and well-distributed mosaic pattern of fiber types, mdx muscles displayed a dramatic structural reorganization marked by a significant increase in the clustering of fiber types, specifically involving glycolytic fast-twitch type II-B fibers (Fig 1H). This marked clustering of fiber types provided morphological evidence of continuous and active denervation-reinnervation cycles occurring early and persistently within the tissue.

**Figure 1:**
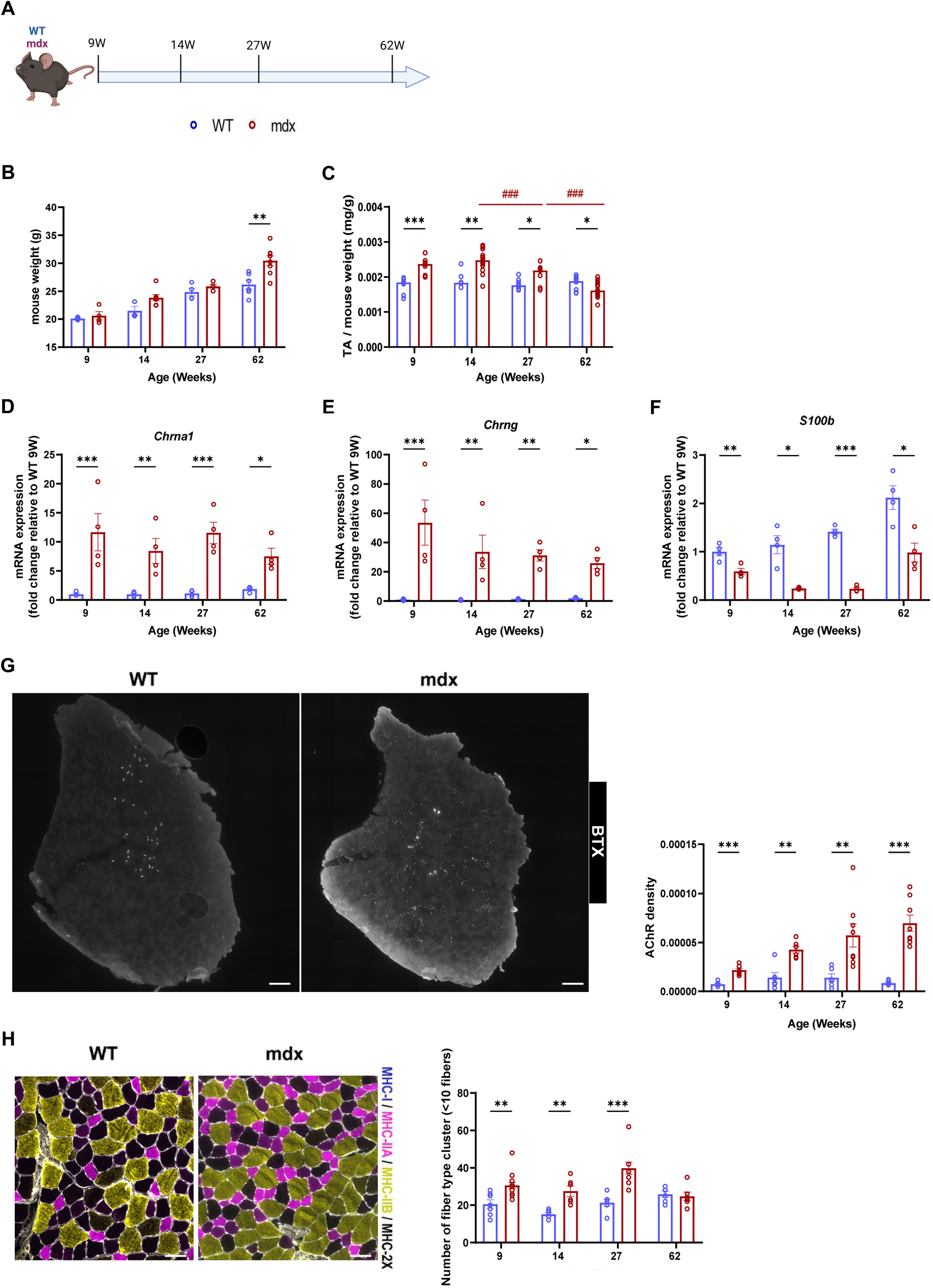
mdx mice display a denervation-like phenotype. (A) Schematic representation of the experimental design. (B) Quantification of body weight (g) in WT (blue) and mdx (red) mice at 9, 14, 27, and 62 weeks of age (N = 4-7) (C) Quantification of Tibialis anterior (TA) mass normalized to body weight (mg/g) in WT (blue) and mdx (red) mice at 9, 14, 27, and 62 weeks (N = 8) (D-F) RT-qPCR analysis of NMJ remodeling markers Chrna1 (D), Chrng (E), and S100b (F) expression in TA muscle at 9, 14, 27, and 62 weeks (N = 4) (G) Representative images of AChR dispersion in WT and mdx TA muscles at 14 weeks, and quantification of AChR density at 9, 14, 27, and 62 weeks (N = 6) (H) Representative images of fiber type grouping in WT and mdx TA muscles at 14 weeks, and quantification at 9, 14, 27, and 62 weeks (N = 6) Data are presented as means ± s.e.m. P-values were calculated using mixed-effects analysis followed by Fisher’s LSD (B,C,G,H), Two-way ANOVA followed by Fisher’s LSD (D,E,F).

Collectively, these findings demonstrate that, despite the pathological muscle hypertrophy, the mdx muscular phenotype is defined by a progressive atrophy and an underlying “denervation-like” state.

### A fast twitch myonuclear denervation program unveils by single nucleus transcriptomics

To decipher the cellular complexity and distinct transcriptional trajectories underlying this neuromuscular uncoupling, we expanded the analysis of our previously established single nuclei RNA sequencing (snRNAseq) dataset of skeletal muscles before and after the necrosis-regeneration cycle from 3 to 9 weeks of age (Fig 2A)(17). Unsupervised clustering and UMAP visualization of the myonuclear landscape (Fig 2B) revealed a distinct temporal dynamic. While completely absent at 3 weeks of age prior to necrosis-regeneration cycle, pathological myonuclear subpopulations uniquely emerged at 9 weeks of age exclusively within mdx muscles. These clusters are characterized by the expression of *Xirp2* and *Scn5a* transcripts linked to physical denervation and synaptic remodeling (Fig 2C-D)(18). To determine the precise cellular lineage of these “denervation-1 & -2” clusters, we executed a pseudo-time trajectory analysis, which demonstrated that these denervated clusters directly segregated from and derived from fast-twitch type II-B myonuclei (Fig 2E). Gene ontology (GO) enrichment analysis of these clusters identified a profound activation of programs orchestrating nervous system development, acetylcholine receptor activity for the “denervation-1” cluster and NMJ maintenance as well as muscle contraction for the “denervation-2” cluster (Fig 2F-G). Furthermore, the comparison of with IIB myonuclei showed an upregulation of many genes where the majority is involved in NMJ maintenance (Fig S2A) GO enrichment analysis identified an activation of neuron projection, axonogenesis and muscle contraction processes (Fig S2B).

**Figure 2:**
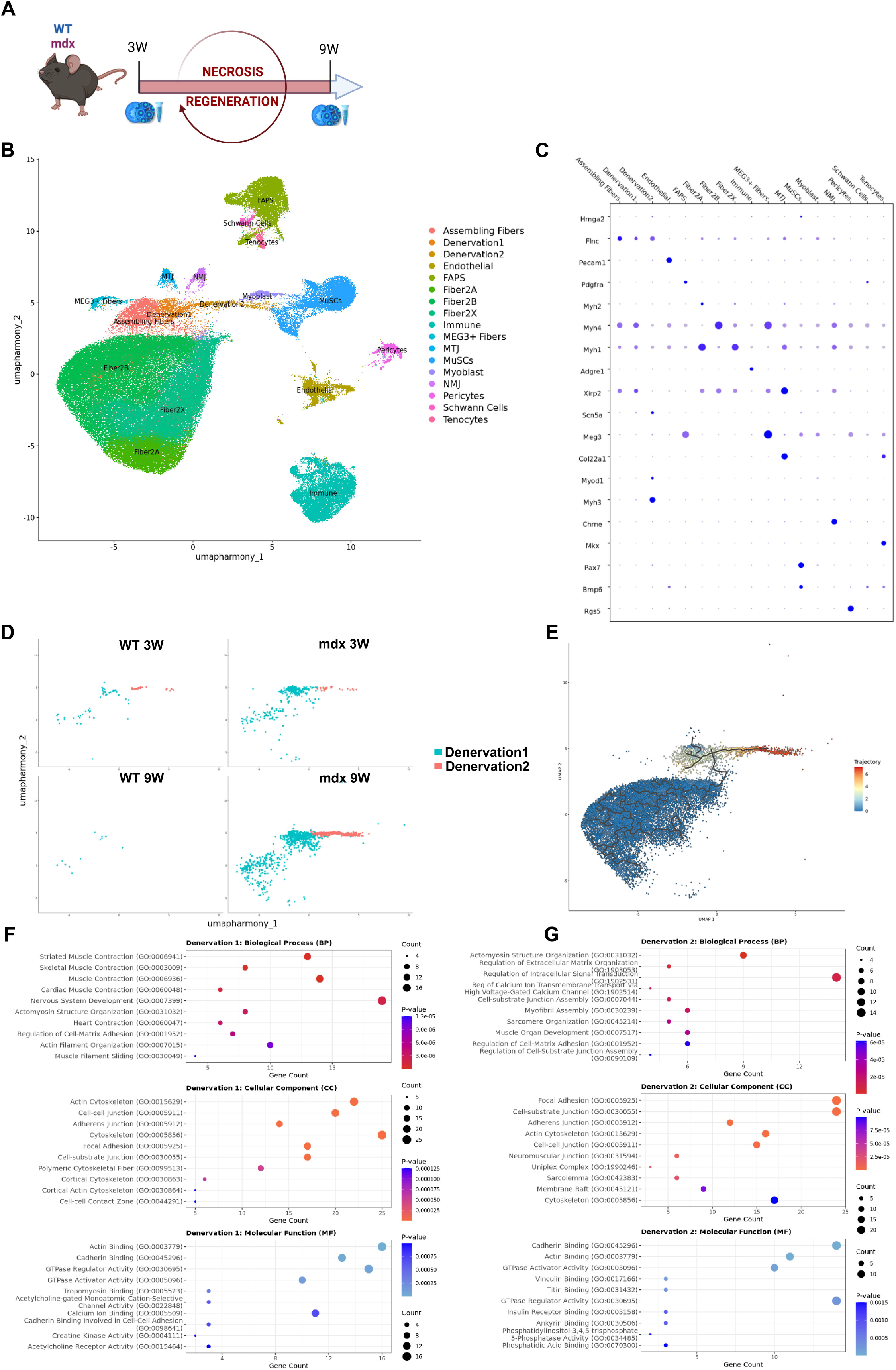
Single-nucleus transcriptomics reveals an active denervation signature in mdx muscles. (A) Schematic representation of the experimental design (B) Unbiased clustering of snRNA-seq data from 3 and 9 week TAs represented on a UMAP plot. Each major cell type is represented by a distinct color (C) Dot plot showing marker genes for major cell types in the TA muscle (D) UMAP plot highlighting the specific presence of Denervation 1 and Denervation 2 clusters in Tas (E) Monocle trajectory analysis demonstrating that Denervation 1 and 2 nuclei originate from type II-B myonuclei (F) Gene Ontology (GO) enrichment analysis for the Denervation 1 cluster (G) Gene Ontology (GO) enrichment analysis for the Denervation 2 cluster Data are presented as means; P-values were calculated with a negative binomial distribution adjusted by Benjamini-Hochberg rank test.

Overall, this high-resolution transcriptomic map provides molecular evidence that a primary, autonomous denervation program is selectively anchored within the II-B myonuclear compartment, establishing a chronic neurogenic pathology.

### Intrinsic primary axonopathy in mdx mice is independent of muscle necrosis

To determine whether the denervation-like signature observed in dystrophic muscle stems from an intrinsic and autonomous alteration of the peripheral nervous system, we characterized the molecular and structural integrity of the sciatic nerve focusing on 14 weeks old mice at the peak of muscle pathological hypertrophy. High-resolution immunofluorescence analysis of cleared teased sciatic nerve fibers revealed major structural anomalies exclusively in mdx mice, characterized by the aberrant accumulation of pathological aggregations of neurofilament and actin along motor axons, confirming a profound disorganization of the axonal cytoskeleton (Fig 3A-B).

**Figure 3:**
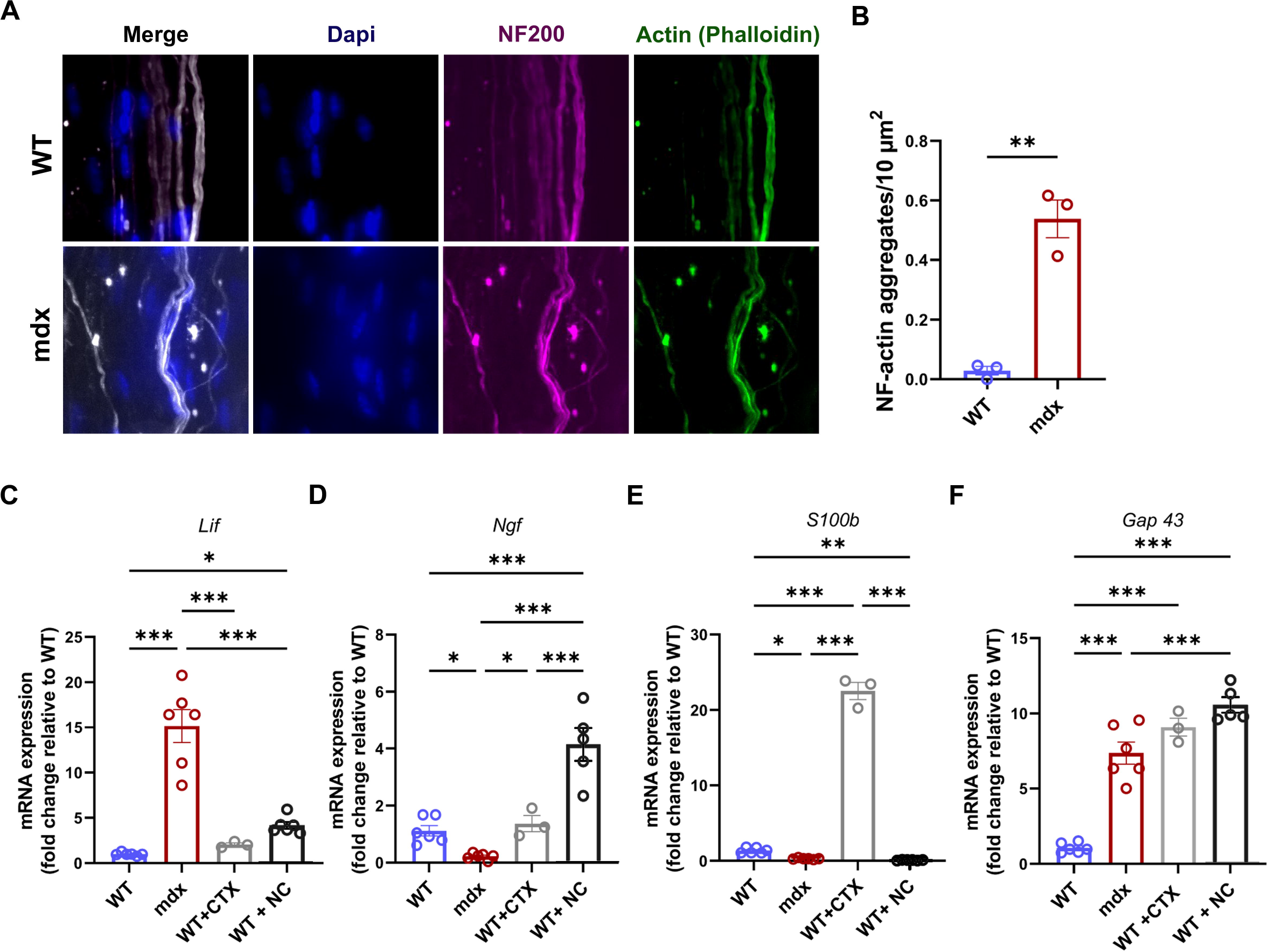
mdx mice exhibit a primary axonopathy. (A) Representative images of whole-mount sciatic nerve immunostaining revealing neurofilament (NF200) and actin (phalloidin) aggregation in WT and mdx mice at 14 weeks (B) Quantification of NF/Actin aggregation (C-F) RT-qPCR analysis of nerve injury and repair transcripts Lif (C), Ngf (D), S100b (E), and Gap43 (F) expression in the sciatic nerve of WT, mdx, cardiotoxin-injected (WT-CTX, 4 days post-injection), and nerve crush (WT-NC, 3 days post-injury) (N = 3-6) Data are presented as means ± s.e.m. P-values were calculated using unpaired t test (B) and Ordinary one-way ANOVA followed by Fisher’s LSD (C,D,E,F)

To establish whether this peripheral axonopathy is a primary autonomous defect or a passive dying-back process secondary to myofiber degeneration, we compared these alterations on sciatic nerves within two models displaying strictly muscle or nerve injury. Specifically, we evaluated WT mice subjected to either muscular cardiotoxin (CTX) injury analyzed 4 days post-injury at the kinetic peak of axonal retraction and a model of mechanical sciatic nerve crush (NC) analyzed 3 days post-injury during acute Wallerian degeneration. Multi- parametric transcriptional profiling revealed a complete molecular decoupling, segregating the mdx nerve phenotype from pure muscle degeneration while mapping it directly to active axonal injury phenotype (Fig 3C-F). The neural distress transcript *Lif* exhibited a massive chronic upregulation in mdx sciatic nerves, significantly surpassing the acute induction observed in NC group, whereas it remained completely unaltered in CTX group (Fig3C). Strikingly, the neurotrophic and regenerative response profiles exposed a fundamental repair defect in the dystrophic nerve. Following NC, WT nerves mounted a robust survival program, characterized by an upregulation of the neurotrophic factor *Ngf expression* (Fig3D). In contrast, mdx nerves failed to initiate this repair drive, exhibiting a severe collapse of *Ngf* transcripts below baseline WT levels (Fig3D). This glio-axonal failure was further substantiated by the SCs markers *S100b,* which collapsed to near-extinction in the mdx and NC groups, yet displayed a massive compensatory upregulation in the CTX group, reflecting a resilient glial response to muscle loss that is entirely lost in dystrophic niche (Fig3E). Lastly, the regeneration associated marker *Gap43* surged across all three conditions, CTX, NC and mdx, compared to WT levels, capturing a universal signature of axonal tip remodeling common to both localized axonal retraction (CTX) and active axonopathy (mdx and NC) (Fig3F).

To figure out whether these acute glio-axonal perturbations represent a transient phenotypical phase or a permanent degenerative state, we performed an extended longitudinal chronological screen. Quantitative RT-qPCR analysis across a wide temporal continuum demonstrated that this chronic neural distress program and axonal structural decay were not restricted to young mice but were robustly sustained as a perennial pathological plateau from 9 weeks of age through 62 weeks (fig. S3A-D).

Together, these specific molecular footprints and long-term kinetic data demonstrate that the peripheral axonopathy in mdx mice possesses its own unique, irreversible pathological identity, characterized by autonomous structural decay and a critical failure of intrinsic neural repair mechanisms independent of muscle dystrophic state.

### The Dp427c dystrophin isoform is expressed in peripheral nerve and accumulates at high mechanical stress regions

To investigate the molecular mechanism underlying the intrinsic primary axonopathy of the mdx nerve and with the hypothesis that dystrophin would exert a direct influence within the peripheral nerve compartment, we assessed the expression and the topographical distribution of dystrophin isoforms within the peripheral nervous system. A robust and selective expression of the transcript of the cortical dystrophin isoform (Dp427c) was detected by RT-PCR specifically in sciatic nerve lysates, whereas it was absent in muscle extracts from WT mice, as expected (Fig 4A). In addition, this transcriptional signature was completely undetectable in mdx nerves, confirming that the *dmd* mutation also abolishes its expression within this tissue (Fig 4A). We next validated this expression at the protein level via western blotting (Fig4B). While WT nerves, as expected, displayed the expected Dp71 and peripheral nerve-specific Dp116 isoforms, we observed an additional band at 427 kDa. This finding confirms the presence of the full-length Dp427c protein within the WT sciatic nerve. As anticipated, this isoform was completely abolished in the mdx sciatic nerve. To precisely map the spatial distribution of the Dp427c protein at a three-dimensional scale without disrupting tissues architecture, we implemented a tissue clearing protocol combining with high resolution 3D confocal imaging on intact sciatic nerves, visualized as maximum intensity projections (Fig 4C). In WT mice, this approach demonstrated that Dp427c intimately follows the cytoskeleton organization of the nerve fiber colocalizing with the neurofilament (NF) and the actin networks. Remarkably, this distribution exhibits a highly stereotyped compartmentalization, as Dp427c accumulation was highly accentuated at specialized structural domains, specifically at the Schmidt-Lanterman incisures (SLI) and at the Nodes of Ranvier (Fig 4C). This topography indicates that Dp427c provides architectural support along the nerve fiber that is strategically reinforced at site of high biomechanical stress(19–21). In mdx mice analyzed at 14 weeks, these networks were entirely absent. The lack of Dp427c deprives the peripheral nerve of this stabilizing support, perfectly coinciding with the disorganization of the NF and the actin networks and the localized structural collapse as demonstrated previously (Fig 3).

**Figure 4:**
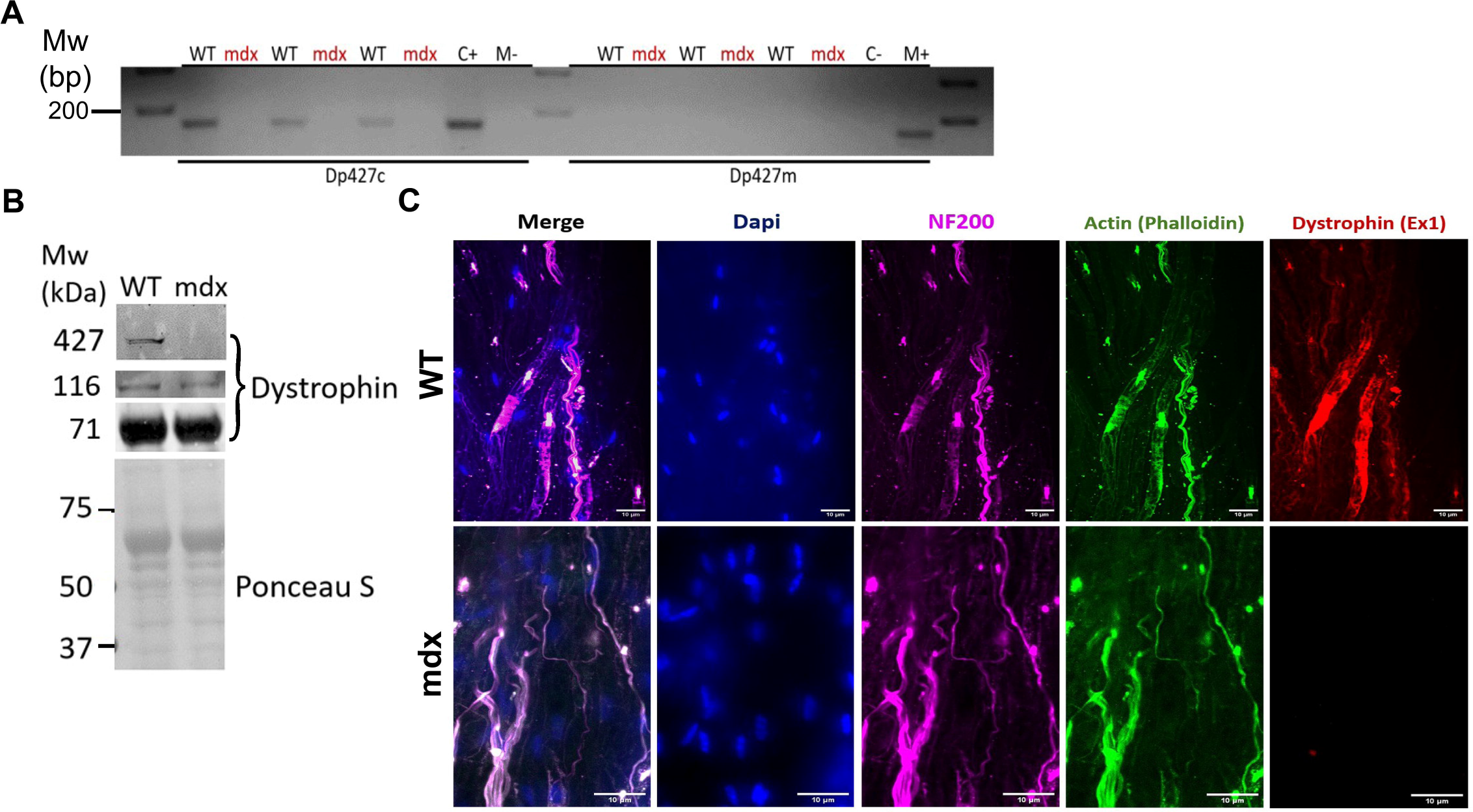
The full-length Dp427c dystrophin isoform is localized at biomechanical stress regions in the healthy peripheral nerve. (A) RT-PCR analysis of Dp427c and Dp427m isoforms in the sciatic nerve of WT and mdx mice. Cortex (C) and Muscle (M) are included as controls of specific isoforms. (B) Western blot analysis of dystrophin isoforms in WT and mdx sciatic nerves (C) Representative images of whole-mount sciatic nerve immunostaining targeting exon 1 of dystrophin

These finding establish the expression of the cortical dystrophin isoform Dp427c within sciatic nerve displaying a specific location and demonstrate that the loss of this protein serves as the primary anchoring defect driving the autonomous axonopathy observed *in vivo*.

### Muscle-restricted microdystrophin expression rescues muscle force but fails to correct neuromuscular transmission deficits

To establish whether restoring sarcolemmal stability is sufficient to rescue the overall integrity of the motor unit, we performed intramuscular injection of an AAV expressing microdystrophin (mdx-µDys) or scrambled sequence (mdx-Scr) under the control of a muscle specific promoter in adult mdx for evaluation at 4 weeks post injection (Fig 5A). First, the specific expression of microdystrophin in muscle was confirmed by mapping vector expression and distribution. Quantification of the viral genome confirmed the presence of the AAV in TA and in the sciatic nerve, demonstrating that the vector had undergone effective retrograde transport (Fig 5B). However, the *microdystrophin* transcript was detected only in TA muscles, while at the sciatic nerve showed no expression of this transcript, thereby confirming that the vector was expressed exclusively in muscle due to the muscle-specific promoter (Fig 5C).

**Figure 5:**
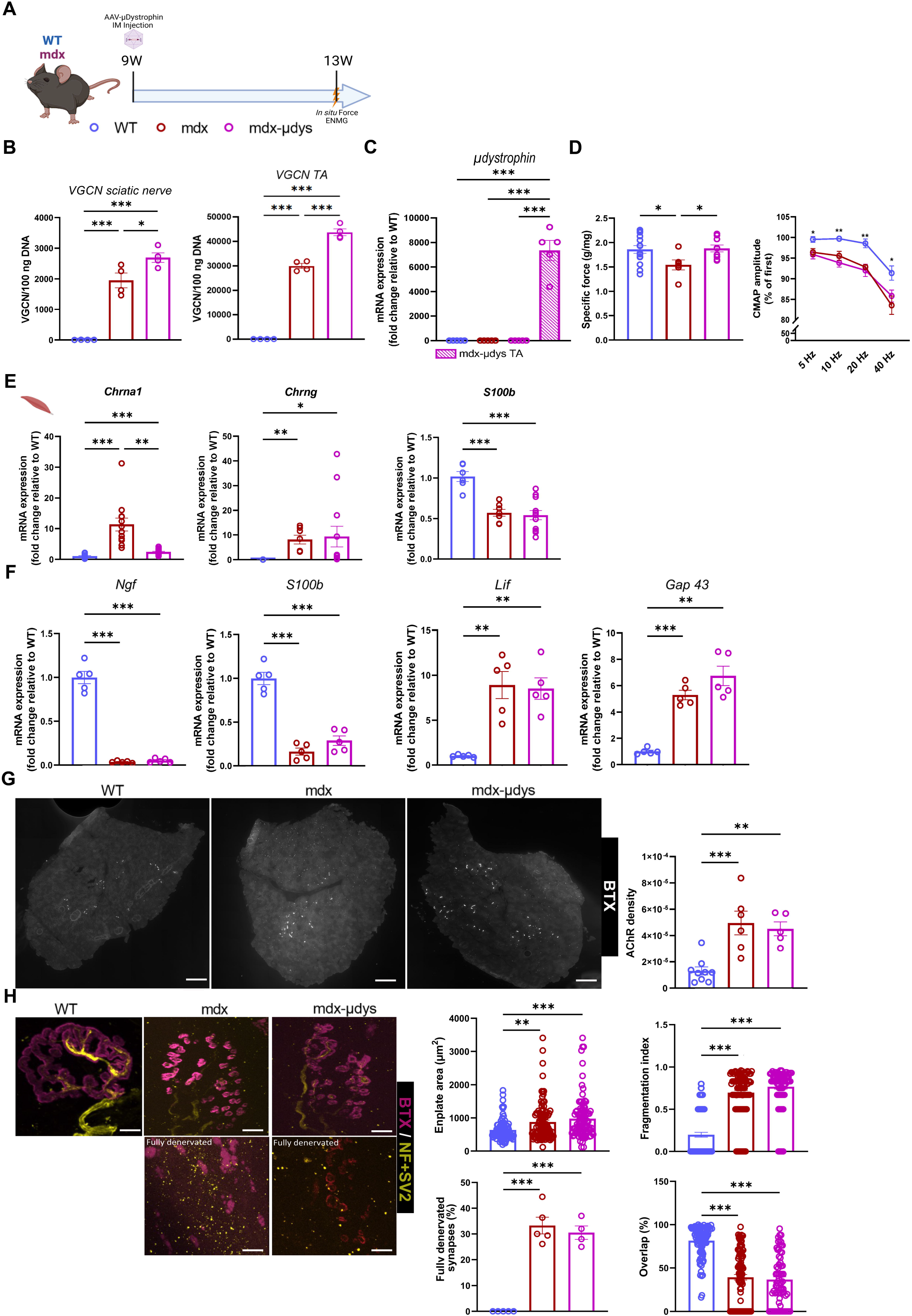
Muscle-restricted microdystrophin expression is not sufficient to restore neurotransmission. (A) Schematic representation of the experimental design (B) Viral genome quantification in the sciatic nerve and TA muscle (C) RT-qPCR analysis of µ-dystrophin expression WT (blue), mdx-scr (red), and mdx-µdys (magenta) (N=6) (D) Specific force measurement and electroneuromyography (ENMG) analysis showing the decrement of the compound muscle action potential (CMAP) in WT (blue), mdx-scr (red), and mdx-µdys (magenta) TA muscles (N = 6-9) (E) RT-qPCR analysis of NMJ remodeling markers Chrna1, Chrng, and S100b expression in TA muscle at 9, 14, 27, and 62 weeks (N = 4) (F) RT-qPCR analysis of nerve transcripts Ngf, S100b, Lif, and Gap43 expression in the sciatic nerve of WT, mdx, and mdx-µdys (N = 3-6) (G) Representative images of AChR dispersion in WT and mdx at 14 weeks, and quantification of AChR density at 9, 14, 27, and 62 weeks (N = 6) (H) Representative images of NMJ morphology from WT, mdx, and mdx-µdys mice analyzed using NMJ-morph. Pre- and post-synaptic regions are stained with NF200+SV2 (Yellow) and α-bungarotoxin (Magenta). Scale bar: 10 µm. Quantification of NMJ hallmarks includes: Endplate area, Fragmentation index, fully denervated synapse, Overlap (N = 100). Data are presented as means ± s.e.m. P-values were calculated using Ordinary one-way ANOVA followed by Fisher’s LSD (B,D,G and H) followed by a Tukey HSD (C), Brown-Forsythe ANOVA followed by Unpaired t with Welch’s correction (E,F) and Kruskal-Wallis followed by Uncorrected Dunn’s (H).z

Biomechanically, the microdystrophin restoration in muscles translated into a complete recovery of specific maximal *in situ* force, validating the efficacy of the gene therapy on the rescue of contractile apparatus activity (Fig 5D). However, *in vivo* electroneuromyography analysis revealed a major functional impairment. Indeed, the amplitude of the compound muscle action potential (CMAP) during repetitive nerve stimulations (5 to 40 Hz) exhibited a severe decrement in mdx mice that was not corrected by microdystrophin treatment (Fig 5D). This critical discrepancy between a functional excitation-contraction coupling and a failing electrophysiological activity indicated a persistent defect specifically localized at the synaptic or presynaptic level.

The molecular origin of this failure was investigated by the assessment of the NMJ remodeling marker expression. Although *Chrna1* transcript was decreased in muscle, likely a secondary benefit of microdystrophin-mediated sarcolemmal stabilization, the other NMJ remodeling marker *Chrng* and *S100b* remained drastically dysregulated (Fig 5E). More importantly, analysis of sciatic nerve integrity revealed that none of the neural integrity markers (*Ngf, S100b, Lif* and *Gap43)* were rescued in mdx-µDys, confirming the persistence of an active state of denervation within the neural compartment (Fig 5F). The consequences of this chronic denervation were corroborated at the muscular level. Pathological fiber type grouping, visualized by an increase in cluster of more than 10 fibers in mdx mice, showed no significant improvement following treatment (Fig S4A). Similarly, the aberrant density and widespread distribution of AChR observed on muscle cross section were not resolved (Fig 5G).

To mechanistically understand this connectivity failure at the sub-cellular level, we analyzed dystrophin topology on isolated muscle fibers. In WT mice, dystrophin colocalized with the sarcolemma and nerve terminals (NF200/SV2 staining) with pronounced enrichment at Node of Ranvier and the motor endplate. In treated mdx mice, while microdystrophin correctly lined the sarcolemma, it was absent from the nerve terminals and completely failed to restore the complex topology of the NMJ (Fig S4B). Final morphometric analysis of the NMJs solidified this observation: microdystrophin was unable to rescue endplate area, AChR fragmentation, the percentage of pre/post-synaptic overlap, and failed to prevent the maintenance of a critical proportion of fully denervated synapses (Fig 5H).

altogether, these results demonstrate that the absence of dystrophin restoration within the neural compartment dooms synaptic architecture, rendering strictly muscle-targeted therapies inefficient.

## Discussion

Historically, DMD has been described by the French neurologist Dr Guillaume Duchenne de Boulogne, as a “pseudo-hypertrophic muscular paralysis”, an initial observation that did not neglect the potential importance of the nervous system(22). Paradoxically, today, DMD is classified and studied under a strictly myo-centric paradigm, defined by sarcolemmal fragility leading to myofiber necrosis. Our study fundamentally challenges this compartmentalized view by demonstrating that the pathology is part of a global collapse of the neuromuscular system. We establish that the mdx dystrophic phenotype, beyond muscle degeneration, is characterized by a state of chronic and autonomous denervation. This alteration is not the mere consequence of passive axonal retraction following the loss of targeted muscle fibers, but results from an intrinsic and primary peripheral axonopathy. At the core of this neural dysfunction is the loss of the cortical dystrophin isoform, Dp427c, whose previously unrecognized expression and crucial architectural role within the peripheral nerve we hereby reveal.

To decode the progression of this dual tissue pathology, our longitudinal study revealed that progressive muscle atrophy is paradoxically masked during the early stages by robust pathological pseudo-hypertrophy due to intense regeneration process and inflammatory cells infiltration(17,23). Beneath this hypertrophy, our snRNAseq data and tissues analysis uncovered an active denervation program, selectively anchored within the myonuclear compartment of fast-twitch type II-B fibers. This chronic instability manifests as an extra- synaptic dispersion of AChRs and fiber type grouping, which are classic morphological signatures of continuous denervation-reinnervation cycles(24). Contrary to the dogma postulating retrograde degeneration (dying-back)(14), the transcriptional profile of the dystrophic sciatic nerve diverged completely from that observed during pure muscle necrosis (CTX model) and overlaps with that of an active axonal injury (NC model). Furthermore, the dystrophic nerve is distinguished by a critical inability to initiate a neural repair program, illustrated by the collapse of neurotrophic transcripts.

Mechanistically, we attributed this axonopathy to the absence of the Dp427c isoform. While our study identifies its critical presence in the peripheral nervous system, Dp427c has been primarily characterized in the central nervous system. In the brain, dystrophin is involved in stabilizing GABA-A receptors and modulating synaptic plasticity, explaining the cognitive and neurodevelopmental disorders observed in some patients(25). We revealed that in a healthy context, this protein intimately follows the cytoskeletal organization of the nerve fiber and accumulates in structural domains subjected to high biomechanical stress, such as SLI and Nodes of Ranvier(19–21). Given that Nodes of Ranvier are exposed, unmyelinated segments essential for impulse propagation, and SLI are dynamic structures conferring plasticity to the myelin sheath, these regions represent potential points of mechanical vulnerability(19,20). The specific enrichment of the Dp427c at these sites suggests a fundamental role in reinforcing the structural architecture of the nerve. In the dystrophic context of mdx, its absence could deprive the peripheral nerve of this stabilizing support, leading to localized structural collapse and the accumulation of pathological actin and neurofilament aggregates(26,27).

Although our results shed new light on the pathophysiology of DMD, the interpretation of our approach must be contextualized against the intrinsic limitations of the mdx mouse model. In this model, the pathology manifests as an acute crisis of necrosis/regeneration followed by phenotypic stabilization with a reduced impact on overall lifespan and motor decline, largely due to strong regenerative capacity and compensation by utrophin(28). These murine-specific traits might mask the true extent of the primary neuropathy observed long-term and limit full appreciation of the neuromuscular failure within a context of progressive exhaustion typical of human patients. Nevertheless, the consistency of the neural distress transcriptomic signature and structural degradation from 9 to 62 weeks in our study attests to the robustness and irreversibility of this neurodegenerative axis, independent of muscle necrosis cycles.

The advent of muscle-targeted microdystrophin gene therapy represents a major preclinical and clinical breakthrough(5). By restricting expression to the muscle via a specific promoter, this approach successfully restores maximal *in situ* contractile force, validating its formidable efficacy in preserving the muscle integrity from necrosis. Nevertheless, evaluating the motor unit as a whole reveals a functional dissociation: although the muscle is mechanically stabilized, the severe decrement in CMAP amplitude persists. This electrophysiological finding suggests that exclusive repair of the sarcolemma, while indispensable, reaches a therapeutic ceiling when it comes to restoring the complex architecture of the NMJ and counteracting synaptic denervation. Because exclusive targeting of the muscle leaves the neural component structurally vulnerable, supporting synaptic transmission and nodal architecture emerges as the next frontier to maximize and sustain the clinical benefits of existing gene therapies. Therapeutically, our work highlights the urgent need to develop combinatorial approaches with multi-tissue targeting. To optimize efficacy while bypassing the potential off-target effects inherent to ubiquitous systemic expression(29), the design of compartmentalized therapeutic strategies, capable of specifically and simultaneously targeting the muscle(5) on one hand and the peripheral nervous system on the other(30), represent a major clinical prospect. Future studies will need to determine whether such cross-targeting is sufficient to stabilize the motor unit in its entirety and resolve the overall functional decline.

In conclusion, this work drives a fundamental paradigm shift, redefining DMD no longer as a strictly myopathic condition, but as an integrated pathology involving motor neurons. This reconceptualization of the motor unit as a whole entity highlights for therapeutic strategies targeting myofiber preservation support by the restauration of peripheral nervous system, an holistic therapy moving from a compartmentalized method to one of targeting the motor unit in its entirely.

## Material and methods

### Animal and Ethical statement

All animal procedures were reviewed and approved by an external ethics committee and the French Ministry of Higher Education and Scientific Research (Project authorization #29280). Experiments were conducted on C57BL/10ScSn-Dmdmdx/J (mdx) and C57BL/6 mice (WT). The animals were housed under SPF conditions under 12h/12h light/dark conditions, with ad libitum access to food and water and cared for following the Directive 2010/63/EU.

### Plasmids and Adeno Associated Virus (AAV) production

pSMD2-Scramble have been generated by direct cloning of a scramble nucleotide sequence in pSMD2 and pSMD2-md1 (microdystrophin) by direct cloning of pAAVSpc512JCMurine [George Dickson: Royal Holloway - University of London (RHUL)], both under spc5-12 promoter. AAV2/9 and AAV2/10 pseudotyped vectors have been prepared by the AAV production facility of the Center of Research in Myology, by transfection in 293 cells as described previously(17). The final viral preparations were kept in phosphate-buffered saline (PBS) solution at −80°C and particle titers (number of viral genomes per ml) was determined by quantitative PCR.

### *In vivo* treatments

Intramuscular administration (mdx): 9-week-old mice were anesthetized using isoflurane (3% induction, 2% maintenance) and injected intramuscularly into the *Tibialis anterior* (TA) with AAV2/10 at 6.10^11^ vg/ml (40µl/TA).

### Electroneuromyography measurements

Electroneuromyography (ENMG) measurements were achieved in TA muscle as previously described(31). During ENMG experiment mouse body temperature was maintained at 37°C with a heating plate. The sciatic nerve was stimulated with series of 10 stimuli at 20 Hz. Compound muscle action potentials (CMAP) were amplified (BioAmp, ADInstruments), acquired with a sampling rate of 100 kHz, filtered with a 5 kHz low-pass and a 1 Hz high-pass filter (Powerlab 8/25, ADInstruments). Peak-to-peak amplitudes were analyzed with LabChart 8 software (ADInstruments). In situ muscle force and ENMG experiments were performed in a blinded manner.

### Force measurement

TA muscle fore contraction was evaluated after sciatic nerve electrical stimulation using supramaximal square-wave pulse of 0.1 ms in duration. Peak absolute force was determined at L0 (the length at which maximal tension was obtained during the tetanus) during isometric contractions in response to nerve electrical stimulation (frequency of 75–150 Hz; train of stimulation of 500 ms). Peak absolute force was normalized against TA muscle weight as an estimate of specific maximal force.

### Tissue harvesting and histology

Following euthanasia (cervical dislocation), skeletal muscle (TA and EDL) and sciatic nerve were rapidly isolate. TA were frozen in liquid nitrogen-cooled isopentane for histological analysis, EDL muscles were fixed in 4% PFA for 1h, while whole sciatic nerves were flash-frozen or fixed in 4% PFA for 5 h. Muscle cryosection (12µm) were generated on a Leica cryostat.

### Immunofluorescence

Muscle cryosections were rehydrated in phosphate-buffered saline (PBS), fixed or not with PFA 4% for 10 min, permeabilized with 0.5% Triton X-100 (Sigma-Aldrich) and blocked in a blocking buffer PBS containing 5% bovine serum albumin, 10% horse serum and 0.2% Triton X-100 for 1 h. Sections were incubated in blocking buffer with primary antibodies overnight at 4°C, washed in PBS, incubated for 1 h with secondary antibodies, thoroughly washed in PBS, incubated with 4′,6′-diamidino-2-phenylindole (DAPI) for nuclear staining for 10 min and mounted in Fluoromount (Southern Biotech). Imaging was performed on a confocal Nikon Ti2 microscope equipped with a motorized stage. Morphometric parameters (cross-sectional area, fiber typing, AchR density) were extracted using deep learning pipelines: Cellpose V3 for cytoplasmic segmentation and StarDist for nuclei, followed by thresholding and morphological quantification in QuPath.

### Neuromuscular Junction (NMJ) and sciatic nerve whole mount staining

EDL muscles and sciatic nerve were fixed respectively for 1h and 5h in 4%PFA after euthanasia. Muscle fiber were isolated by mechanical dissection, quenched in 0.1M Glycine-PBS and permeabilized. Samples incubated at 4°C overnight with Neurofilament antibody diluted in blocking solution. Isolated fibers were washed several times in 0.1% Triton-PBS, then incubated at 4°C overnight with a secondary antibody and α-Bungarotoxin-Alexa Fluor™ conjugated 594. Whole mount staining of the sciatic nerve follows a similar permeabilization and immunolabeling protocol to assess structural integrity. Prior to mounting, immunolabeled sciatic nerves were subjected to a tissue clearing process using successive glycerol bath (25%, 50% and 75%), with each incubation lasting 24h. Isolated fibers and sciatic nerves were then mounted on glass slide with VECTASHIELD® Antifade Mounting Medium (Eurobio Scientific) and kept at 4°C until image acquisition.

For NMJ morphology, images were acquired with a confocal microscope (Zeiss 63X objective) and edited with Zeiss Zen Lite 3.7 software. The same laser power and parameter setting were used throughout to ensure comparability. The images presented are single projected images obtained by overlaying sets of collected z-stacks. NMJ morphometric analysis was performed on confocal z-stack projections of individual NMJs by using ImageJ-based workflow adapted from NMJ-morph. At least 100 NMJs have been analysed per condition. NMJ morphometric analysis was performed in a blinded manner by the same investigator.

For Sciatic nerve whole mount staining, Imaging was performed on a confocal Nikon Ti2 microscope equipped with a motorized stage. (100X objective).

### Immunoblotting

Cryosections from frozen TA muscles were homogenized with a dounce homogenizer in a cell lysis buffer (Cell Signaling) supplemented with phosphatase inhibitor cocktail (Roche). Lysates were centrifuged for 5 min at 1500g and protein concentration was determined in supernatant with Bradford method using Protein Assay Dye Reagent (Bio-Rad). Proteins were denatured at 95°C for 5 min with Laemmli buffer and β-mercaptoethanol (10% v/v) and then separated by electrophoresis (Nu-PAGE 4–12% Bis-Tris gel; Thermofisher Scientific) and transferred to nitrocellulose membranes (GE Healthcare). Membranes were blocked with 5% skimmed milk diluted in Tris Buffered Saline (TBS)-0.1% Tween (TBS-T) and incubated at 4°C overnight with primary antibodies. After washes in TBS-T, membranes were incubated with secondary antibodies conjugated to a fluorophore (Bio-Rad). Images were acquired with ChemiDoc™ MP (Bio-Rad) and band intensity was quantified using Image Lab software (Bio-Rad).

### Nucleic acid extraction RT-qPCR and viral genome quantification

Total RNA was isolated from mouse muscles using TRizol (Life Technologies) and Direct-zol RNA Miniprep (Zymo research). From PFA-fixed spinal cord using RNeasy FFPE Kit (Qiagen) according to manufacturer protocol. Complementary DNA was generated with Superscript II Reverse transcriptase (Life Technologies), and analyzed by real time qPCR. Real-time qPCRs were performed on QuantStudio 3 and 7 Pro Real-Time PCR System (Applied Biosystems) using Power SybrGreen PCR MasterMix (Applied Biosystems). All data were analysed using the 2-ΔΔCT method and normalized to P0 (mouse acidic ribosomal phosphoprotein P0) mRNA expression. Classic end-point PCR products were additionally resolved on standard 1.5% agarose gel. For AAV genome quantification, DNA was extracted using the PureGene tissue core kit (Qiagen). Viral genomes were quantified by duplex qPCR targeting the transgene promoter, normalized to total genomic DNA.

### Single nuclei RNA sequencing

Nuclei from cryopreserved TA muscle were isolated using 10x Genomics Chromium Kits, labelled (Acridine Orange/Propidium Iodide) and counted (Luna FL). Libraries were generated utilizing the Chromium Next GEM single 3’ Kit version 3.1 and sequenced on a NextSeq 2000. Data were pre-processed using SoupX, demultiplexed and mapped using Cell Ranger. Downstream analysis was conducted in Seurat v5, filtering out low-quality nuclei. Clustering utilizing specific marker including Xirp2 and Scn5a for denervation signatures. Pseudotime trajectory inference were generated using Monocle3. The raw dataset is available under GEO accession number GSE309516(17).

### Statistics analyses

For comparison between two groups, data were tested for normality using a Shapiro–Wilk test and for homoscedasticity using a Bartlett test followed by parametric (two-tailed paired, unpaired Student’s t-tests) or non-parametric test (Mann-Whitney) to calculate P values (as detailed in the figure legends). For comparison among more than two groups, according to normality ordinary one-way ANOVA or Kruskal-Wallis tests were performed. According to homoscedasticity, Brown-Forsythe ANOVA were performed. When it was necessary, two-way ANOVA tests were performed to compare between groups (as detailed in the figure legends). All ANOVA and Kruskal-Wallis were followed by appropriated post-hoc tests. All statistical analyses were performed with GraphPad Prism 9 software. Statistical significance was set at P < 0.05 and all bar graphs presented are means ± SEM. Asterisks indicate significant differences (*P < 0.05; **P < 0.01; ***P < 0.001) between groups, hashtags indicate significant differences (#P < 0.05; ##P < 0.01; ###P < 0.001) in same group between different timepoint according to statistical analysis performed.

## Acknowledgments

We would like to thank UMS28 animal facility for animal care and technical support, the MyoImage Platform of the Center of Research in Myology (Institut de Myologie, Paris, France) for image analysis and We thank Pierre Meunier and Sofia Benkhelifa-Ziyyat from the MyoVector technical platform of the Centre of Research in Myology-UMRS974 (Paris, France) for AAV production.

## Author contributions

AB and FPR conceived and designed the study. AB, MG, AF, LS, PM, LM, JP, ML and CG performed the experiments. AB, SB, CP, CG and FPR analyzed the data. AB and LG analyzed bioinformatic data. AB and FPR wrote the original draft. CG and SF writing review and editing. FPR supervised the project.

## Data, code, and materials availability

All data are available in the main text. Codes were used according to developer tutorial and available in the “Material and methods” section.

