## Supplemental Material for "Axonopathy in Duchenne Muscular Dystrophy limits microdystrophin gene therapy efficacy"

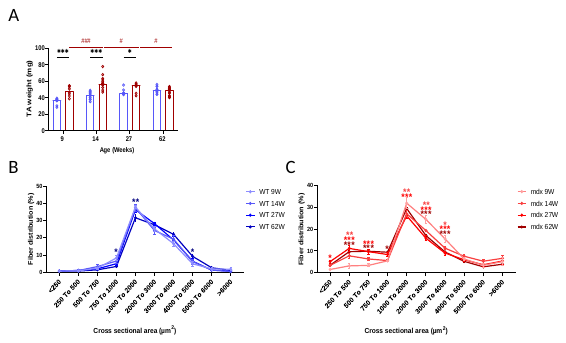
**Supplemental Material**

**Figure S1 : mdx mice display progressive muscle atrophy**

**A:** Absolute TA mass evolution over time in WT (blue), mdx (red)

**B:** Cross-sectional area (CSA) distribution in WT mice over time

**C:** Cross-sectional area (CSA) distribution in mdx mice over time

Data are presented as means ± SEM. P-values were calculated using mixed-effects analysis followed by Fisher’s LSD (A,C), Two-way ANOVA followed by Fisher’s LSD (B). Asterisks indicate significant differences compared to WT 9W (B) and to mdx 9W (C)

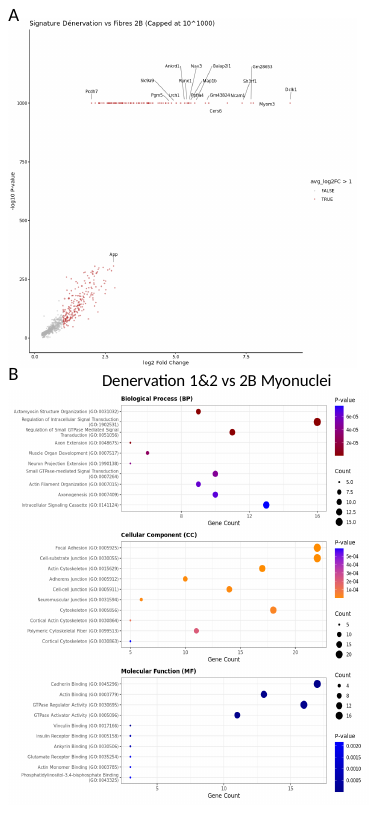

**Figure S2: Denervation clusters overexpress NMJ transcripts**

**A:** Volcano plot of differentially expressed genes between Denervation 1 and 2 clusters and type II-B fiber myonuclei

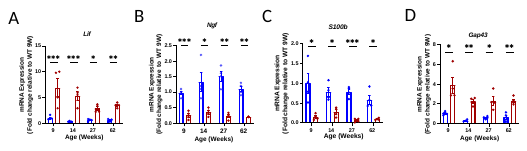
**B:** Gene Ontology (GO) analysis comparing Denervation 1 and 2 clusters to type II-B fiber myonuclei.

**Figure S3: mdx axonopathy persists over time**

A-D: RT-qPCR analysis of nerve transcripts *Lif (*A)*, Ngf* (B)*, S100b* (C)*, Gap43*(D) expression in the sciatic nerve of WT and mdx mice (N = 3-6)

Data are presented as means ± SEM. P-values were calculated using Two-way ANOVA followed by Fisher’s LSD (A,B,C,D)

**Figure S4: Muscle-restricted microdystrophin does not rescue the denervation-like phenotype**

**A:** Representative images of fiber type grouping in WT, mdx, and mdx-µdys muscles

**B:** Representative images of dystrophin expression at the NMJ

Data are presented as means ± SEM. P-values were calculated using Ordinary one-way ANOVA followed by Fisher’s LSD (A).

**Listing of primers**

| **Gene** | **Forward** | **Reverse** |
| --- | --- | --- |
| ***Chrna1*** | AAGCTACTGTGAGATCATCGTCAC | TGACGAAGTGGTAGGTGATGTCCA |
| ***Chrng*** | GCTCAGCTGCAAGTTGATCTC | CCTCCTGCTCCATCTCTGTC |
| ***Dp427c*** | GGCATGGAAGATGAAAGAGAAGA | GGCAGTTTTTGCCCTGCTAAGG |
| ***Dp427m*** | GGACTGTTATGAAAGAGAAGATGTTCA | TGGCAGTTTTTGCCCTGCTAAG |
| ***Gap43*** | CCTGCTGCTGTCACTGATGCTG | CCTGCTGCTGTCACTGATGCTG |
| ***Lif*** | GTCTTGGCCGCAGGGATTG | GCACAGGTGGCATTTACAGG |
| ***µ-dystrophin*** | CGTGATGGAGACCGTGACC | CCTTGATCACTTCGGCCTGT |
| ***Ngf*** | GCAGTGAGGTGCATAGCGTA | CTGTGTCAAGGGAATGCTGA |
| ***S100β*** | CTTCCTGGAGGAAATCAAGGAG | CTCATGTTCAAAGAACTCATGGC |

**Listing of primary antibodies**

| **Name /Target** | **Source** | **Reference** | **Species** | **Application** | **Dilution** |
| --- | --- | --- | --- | --- | --- |
| Dystrophin Ex1 | DSHB | MANEX 1B | Mouse | IF | 1/50 |
| Dystrophin (C-ter) | Leica Biosystems | NCL-Dys2 | Mouse | WB | 1/100 |
| MYHI | DSHB | BA-D5-s | Mouse | IF | 1/10 |
| MYHIIa | DSHB | SC-71s | Mouse | IF | 1/10 |
| MYHIIb | DSHB | BF-F3-s | Mouse | IF | 1/10 |
| Neurofilament | Millipore | AB9568 | rabbit | IF | 1/500 |
| Phalloidin CruzFluor 555 | Santa Cruz | sc-363794 |  | IF | 1/1000 |
| SV2-c | DSHB |  | Mouse | IF | 1/500 |
| α-Bungarotoxin-Alexa Fluor^TM^ 594 | Thermofisher | B13423 | B. multicinctus | IF | 1/500 |
| Laminin α2 | Santa Cruz | sc-59854 | Rat | IF | 1/50 |
